# SMAP: exploiting high-throughput sequencing data of patient derived xenografts

**DOI:** 10.1101/440008

**Authors:** Yuna Blum, Aurélien de Reyniès, Nelson Dusetti, Juan Iovanna, Laetitia Marisa, Rémy Nicolle

## Abstract

**Background:** Patient-derived xenograft is the model of reference in oncology for drug response analyses. Xenografts samples have the specificity to be composed of cells from both the graft and the host species. Sequencing analysis of xenograft samples therefore requires specific processing methods to properly reconstruct genomic profiles of both the host and graft compartments.

**Results:** We propose a novel xenograft sequencing process pipeline termed SMAP for Simultaneous mapping. SMAP integrates the distinction of host and graft sequencing reads to the mapping process by simultaneously aligning to both genome references. We show that SMAP increases accuracy of species-assignment while reducing the number of discarded ambiguous reads compared to other existing methods. Moreover, SMAP includes a module called SMAP-fuz to improve the detection of chimeric transcript fusion in xenograft RNAseq data. Finally, we apply SMAP on a real dataset and show the relevance of pathway and cell population analysis of the tumoral and stromal compartments.

**Conclusions:** In high-throughput sequencing analysis of xenografts, our results show that: i. the use of *ad hoc* sequence processing methods is essential, ii. high sequence homology does not introduce a significant bias when proper methods are used and iii. the detection of fusion transcripts can be improved using our approach. SMAP is available on GitHub: cit-bioinfo.github.io/SMAP.

## Background

In oncology, Patient-derived xenografts (PDX) is to date the closest available model to the human disease. Despite having obvious a *priori* limitations, *e.g.* an altered microenvironment with species divergence, their resemblance with primary tumors is manifold higher than in vitro cell culture. PDX are increasingly used for many aspects of clinically relevant cancer research programs such as drug discovery or identification of drug resistance mechanisms [1, 2]. Among the most promising aspect of PDX models is the possibility to test novel treatments while generating large-scale molecular profiles on the same tumor sample. The resulting data can then be used to identify predictive biomarkers. Deep sequencing is the method of choice to generate exhaustive molecular profiles. However, sequencing PDX samples has a major pitfall: it necessarily involves sequencing RNA or DNA from both human grafted cells and host cells, generally mice. Although this might appear as unwanted and a waste of costly sequencing reads, it is on the contrary a unique opportunity to accurately capture simultaneously molecular signals from the tumor and the stroma [3]. Indeed, the cellular composition and function of the tumor microenvironment has a major impact on key aspects of the disease such as tumor initiation, growth, metastasis or therapeutic resistance [4, 5, 6]. This unique potential of studying the microenvironment of tumors in an *in vivo* model defines a particularly challenging task of simultaneously analyzing a mixture of sequencing reads from two distinct but phylogenetically-close species.

Several bioinformatics approaches have been proposed and used to process sequencing data from the mix of host and graft cells. The simplest one consists in independently mapping the entire set of xenograft reads to the human genome or the mouse genome only and quantify the transcriptomes from thereon. We termed this approach “independent mapping”. Intuitively, this approach may introduce strong quantification bias in homologous genes. The first methodology specifically designed to process xenograft sequencing called Xenome [7], uses an interesting k-mer dictionary approach to assess the species from which each read originates. It is based on the unexpectedly small number of common k-mers in the transcriptomes of human and mice. Xenome can simply be used to classify and separate reads prior to their mapping. Another approach, S3 for Species-Specific Sequencing [8], was recently used in breast cancer to assign human and mouse read origin from xenograft transcriptome profiles. S3 relies on post-mapping processing by filtering out any reads that mapped to the murine genome with a predefined mapping score difference. Similarly to S3, an additional approach named Disambiguate proposes to align reads to both species independently and assign reads to the species with higher quality alignments [9].

In this study, we propose a novel approach termed SMAP for Simultaneous Mapping. SMAP uses the mapping step to select the best matching locus for each gene in either genome.

## Results

### Overview of SMAP: Simultaneous Mapping for Patient-derived xenografts

SMAP is a mapper-agnostic method that runs in three steps to assign each sequencing read (or read pair) to the most relevant species (see Fig 1).

First, chimeric genome and transcriptome are constructed by the concatenation of the host and graft species genomes. Then, a conventional alignment pipeline processes the unsorted sequencing reads using the chimeric genome, thereby mapping all sequences simultaneously on both genomes. Finally, aligned reads are separated based on which part of the chimeric genome they are best aligned to, host or graft, and all reads with an identical alignment score on both genomes are removed. The output of SMAP is a strictly distinct pair of aligned read files (BAM) or gene count matrices, one for the graft and one for the host. Common analysis can then be applied separately such as mutation calling, copy number analysis or gene expression and pathway analysis. As immunocompromised mice are the most widespread chassis for PDX, we will hereafter refer to the host species as murine and the graft as human.

### Overview of the SMAP-fuz methodology

Reliable detection of chimeric transcripts, an aberrant genetic event of gene fusion with high pathogenic potential especially in cancer, is proposed here to also take advantage of the specificity of PDX sequencing. Fusion detection is a challenging task and a large number of methods have been devised to perform it from RNAseq data (reviewed in [10, 11]).

**Figure 1.**
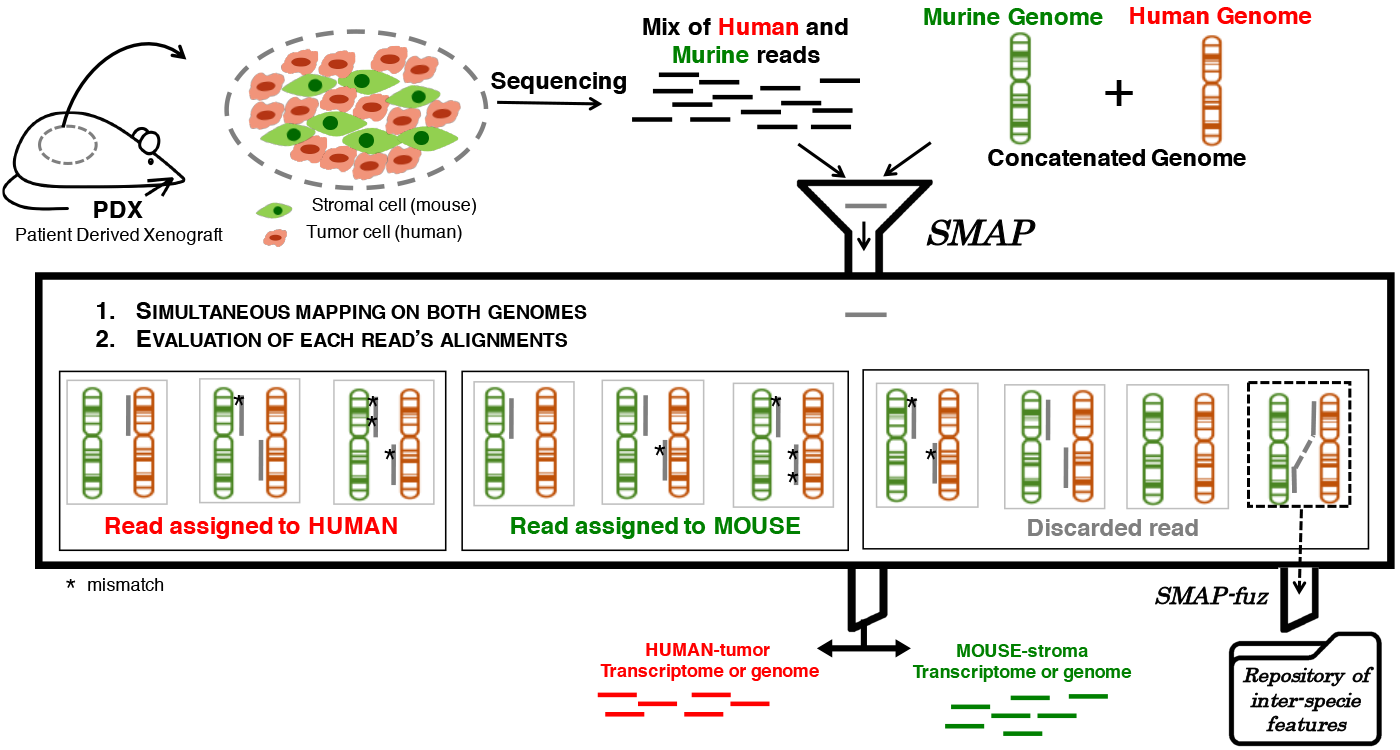
Schematic representation of SMAP. SMAP is designed to process high-throughput sequencing reads from PDX samples which are considered as a mix of tumor cells of the grafted species (usually human) and of stromal cells from the host species (usually mouse). Each sequencing read is processed by a mapping pipeline which uses both genomes (and/or transcriptomes) of both species. Reads are simultaneously mapped to both genomes and only those for which the first and only best alignment is made on one of the two species genome are kept and assigned to their respective species. Typical mapping cases ending in the assignment of a read to the human, murine genome or neither are shown. In the special case of a read partially mapping to two distinct chromosomes from both species, the alignment is stored for use in subsequent analysis of fusion transcript detection.

The problem to tackle is the removal of the excessive rate of spurious fusions that are due to local mapping errors. The core idea relies on the assumption that no chimeric transcript can result from the fusion of a mouse gene with a human gene. Therefore, the characteristics - in terms of junction read and spanning fragment counts - of these false fusions are used to define a null probability distribution and compute a statistics for any potential human-human fusion. The SMAP fusion approach is described in Fig 2 in which it is applied to STAR-Fusion output. However, the process is independent of the fusion detection algorithm as it only requires metrics to build the null probability distribution.

Null distributions of junction read count and spanning fragment count were estimated by fitting a Negative Binomial distribution on their empirical distribution for interspecies fusions - fusions between a human gene and a mouse gene - which were considered as spurious fusions:

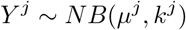

and

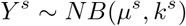

where *μ*^*j*^ and *μ*^*s*^ are the mean of junction and spanning read counts distributions respectively and and *k*^*j*^ and *k*^*s*^ are the dispersion parameter of junction and spanning read counts distributions respectively.

**Figure 2.**
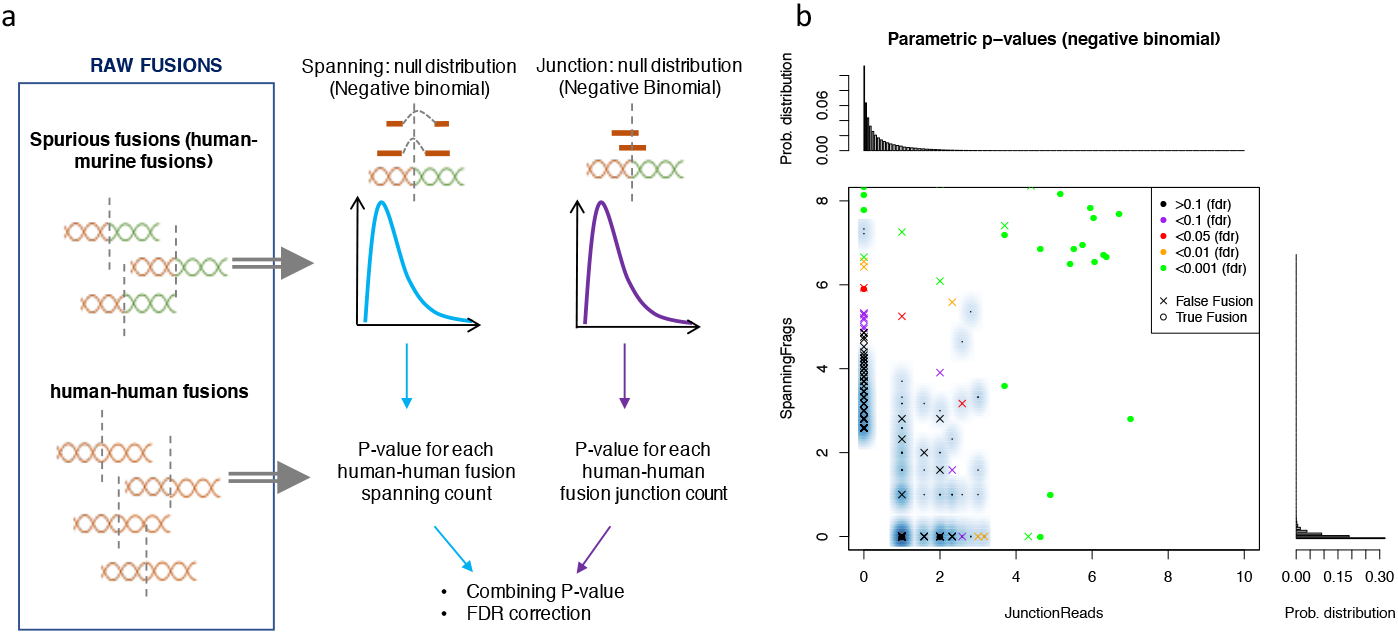
SMAP-fuz approach. **a**. Overall SMAP-fuz approach based on the repository of inter-species fusion transcripts, *i.e.* detected fusions between human and murine genes. A null distribution is built from the available fusion characteristics, here the number of spanning fragments and of junction reads, to model the background noise of fusion detection. These distributions are used to then score the human-only fusion transcripts detected. **b**. Illustrative example of SMAP-fuz results on simulated data. Plot shows observed spanning and junction counts. Dots correspond to fusions and are colored according to their corresponding FDR P-value computed by SMAP-fuz. Spurious fusions (inter-species fusions) are represented by a smoothed color density scatterplot.

The variance of the Negative Binomial distribution is 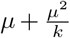 in this parametrization. For each human fusion, p-values for junction reads and spanning fragments were calculated giving these distributions. We used Fisher’s Method [12] to combine both p-values as follows:

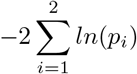

Under null hypothesis (equivalent to all single null hypotheses true) this statistics follows a Chi-squared distribution with 4 degrees of freedom. Corresponding P-values were calculated for each human fusion and were corrected for multiple testing using FDR method.

Null distributions of junction read count and spanning fragment count were estimated by fitting a Negative Binomial distribution on their empirical distribution for interspecies fusions - fusions between a human gene and a mouse gene - which were considered as spurious fusions:

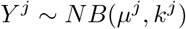

and

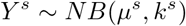

where *μ*^*j*^ and *μ*^*s*^ are the mean of junction and spanning read counts distributions respectively and *k*^*j*^ and *k*^*s*^ are the dispersion parameter of junction and spanning read counts distributions respectively.

The variance of the Negative Binomial distribution is 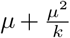 in this parametriza-tion. For each human fusion, p-values for junction reads and spanning fragments were calculated giving these distributions. We used Fisher’s Method [12] to combine both p-values as follows:

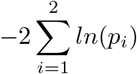

Under null hypothesis (equivalent to all single null hypotheses true) this statistics follows a Chi-squared distribution with 4 degrees of freedom. Corresponding P-values were calculated for each human fusion and were corrected for multiple testing using FDR method.

### Mapping and species assignment performance

In order to assess the proposed method, we simulated xenograft RNAseq data with a mix of human and mouse reads using diverse settings of sequencing errors, mutations rate and read length (see table 1). The purpose of this simulation analysis is to compare various approaches for Xenograft sequencing data processing and to compare their ability to assign each read to the correct species.

**Table 1.** Sample table title. This is where the description of the table should go.

|  | Species | Read length (bp) | Mutation rate | Error rate |
| --- | --- | --- | --- | --- |
| Ideal | human | 50 | 0 | 0 |
|  | mouse | 50 | 0 | 0 |
| Mutations | human | 50 | 0.01 | 0 |
|  | mouse | 50 | 0.001 | 0 |
| Mutations and errors | human | 50 | 0.01 | 0.001 |
|  | mouse | 50 | 0.001 | 0.001 |
| Ideal | human | 100 | 0 | 0 |
|  | mouse | 100 | 0 | 0 |
| Mutations | human | 100 | 0.01 | 0 |
|  | mouse | 100 | 0.001 | 0 |
| Mutations and errors | human | 100 | 0.01 | 0.001 |
|  | mouse | 100 | 0.001 | 0.001 |

The simulated datasets were processed using five different strategies: naively, by independent mapping the entire sequencing data to the human or mouse genomes and using SMAP, Xenome, S3 or Disambiguate. Figure 3 shows the rate of human reads correctly assigned to the human genome (True Positives), the rate of lost human reads that were unmapped or considered as ambiguous (False negatives) and the rate of murine reads assigned incorrectly to the human genome (False Positives). Supplementary figure 1 shows the same results for the murine genome.

The results demonstrate that mapping a mix of human and murine sequences directly onto the human genome results in a large number of false positive murine sequence alignments to the human genome. Disambiguate exhibits as well poor results in terms of false positive rate showing that its criteria based on quality alignment score used to distinguish species is not sufficient to correctly assign reads.

SMAP and S3 as well as to some extent Xenome align virtually no murine sequences to the human genome, approaching an ideal specificity. However, Xenome and in particular S3 systematically discards more human sequences by considering them as ambiguous. Overall, these results show that specific methods are necessary to process PDX sequencing data to avoid significant bias in downstream analysis.

**Figure 3.**
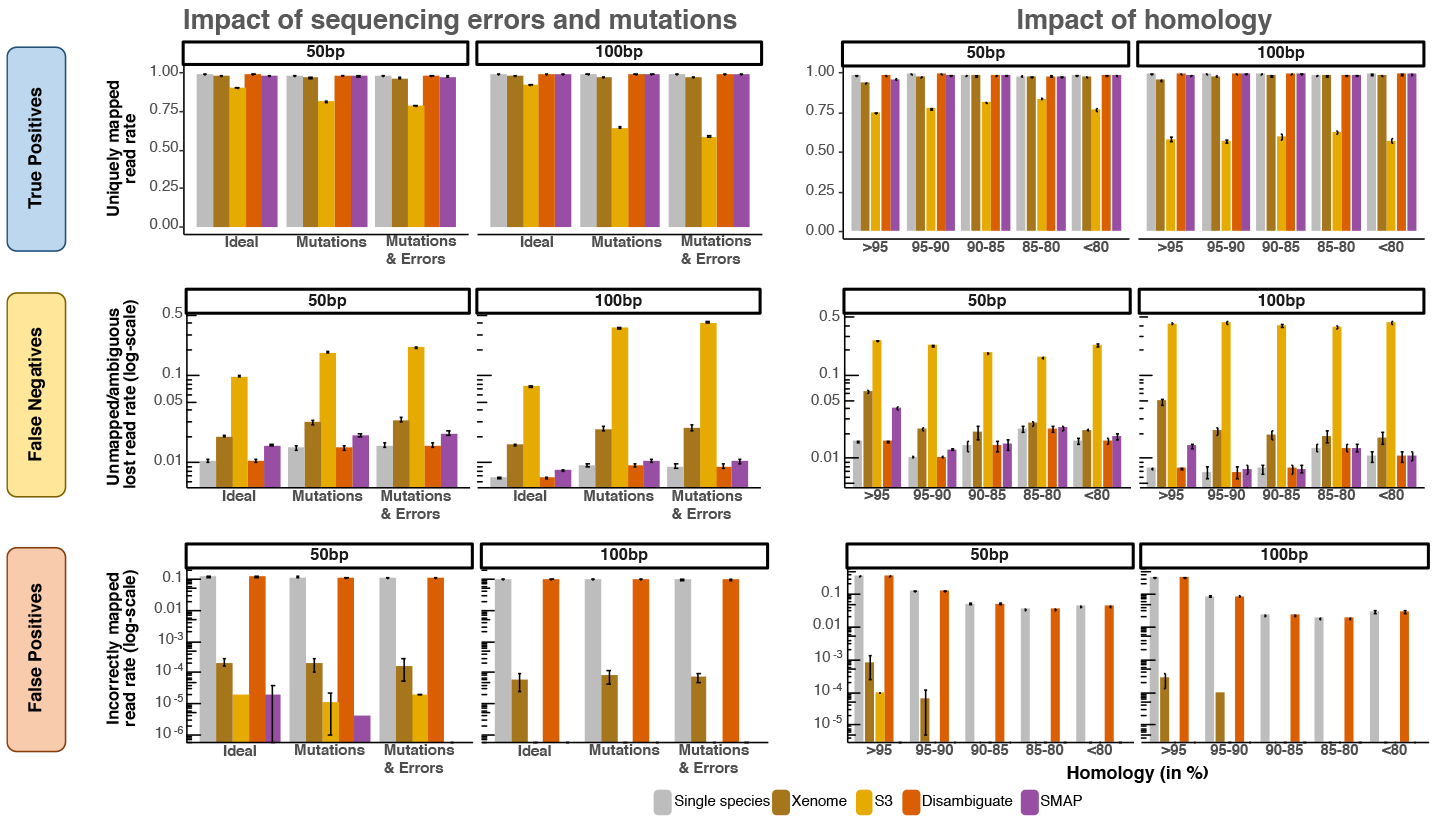
Performance of reads mapping to the human genome. Barplots of the true positive (**a** and (**d**), false negatives (**b** and (**e**) and false positive (**c** and (**f**) rates obtained by the different alignment strategies of mixed human and murine reads. Bars represent the mean and error bars the standard deviation of five replicate simulation experiments. **a.** Proportions of the human reads mapped to the human genome (True Positive), **b**. Proportions of the lost human reads unmapped or ambiguous (False negatives) and **c.** Proportions of the murine reads mapped to the human genome (False Positive) in various settings of read simulation parameters (see table 1). **d.** Proportions of the human reads mapped to the human genome (True Positive) **e.** Proportions of the lost human reads unmapped or ambiguous (False negatives) and **f.** Proportions of the murine reads mapped to the human genome (False Positive) in simulations on sets of genes with various levels of homology between the human and murine orthologs.

It is expected that there is a high level of discrepancies between the reference genomes and the sequencing reads, which can either originate from sequencing errors or genetic variances (*e.g.* polymorphism, mutations), may degrade the capability of sequence processing pipeline to assign reads to their correct species. As reported in figure 3, three of the tested pipelines suffer only moderately from these divergences with correct assignment rates dropping in a realistic setting of less than 1% for independent mapping (100bp: 2.4‰, 50 bp: 5.3‰), Xenome (100bp: 9‰, 50 bp: 1.1%) or SMAP (100 bp: 2.3‰, 50 bp: 5.9‰) as compared to an ideal setting. S3, which is overall much less performant, is greatly impacted by the number of discrepancies between the sequencing reads and the genome with 33.7% drop in the rate of correct assignment between the ideal and mutation and error setting at 100 bp (11.4% at 50bp). On the other hand, errors in species-assignment are not impacted by discrepancies between sequencing reads and the genome. Altogether, sequencing longer reads (*i.e.* 100 bp compared to 50 bp) increases the rate of correct and decreases the rate of incorrect species-assignment for every method, except S3. The exceeding stringency of S3 could be explained by a higher absolute number of mapping errors in longer reads in a constant error rate setting. Likewise, S3 gave better results on the murine genome, as mouse reads were simulated with lower mutation rate (Supplementary figure 1). Overall, the use of SMAP results in better read assignment with a significantly lower rejection rate and a virtually null rate of false species-assignment even in highly mutated samples.

### Estimating the impact of inter-species gene homology

The homology between the coding sequences of Humans and Mice is the prime difficulty to overcome in PDX genomic studies. Sequencing reads from human genes which have a highly homologous murine ortholog are expected to be more frequently assigned to the wrong species genome.

In order to quantify the impact of homology on xenograft sequencing, reads were simulated specifically on subsets of orthologous genes with variable levels of sequence similarity. In figure 3, we show that the species-assignment errors of sub-optimal methods are highly correlated with the level of sequence similarity. For instance, naively mapping the mix of human/murine PDX reads to the human genome as well as Disambiguate approach result in more than 30% of the murine reads incorrectly assigned to the human genome in highly homologous genes (more than 95% similarity). This rate drops significantly to 12% and below 10% in orthologous genes with lower levels of homology. Homology has less impact on the number of false positives in other methods with only Xenome misclassifying less than 1 per thousand reads for highly homologous genes.

Overall, SMAP systematically outperforms other method by assigning more reads to their correct genome while maintaining a low near-null false positive rate.

### Evaluation of SMAP-fuz

In order to quantify the impact of SMAP on fusion detection, the processing pipeline was applied to simulated transcriptomes with known fusion events. Figure 4 reports several accuracy metrics comparing the detection of fusion with or without SMAP-fuz. The proposed approach does not remove true fusions while discarding a large number of wrongly detected fusions (80% for 50bp reads, 65% for 100bp reads). Overall, using SMAP-fuz for fusion detection results in an extensive improvement in false discovery (as measured by the positive predictive value PPV) with virtually no trade-off in sensitivity.

### Application to pancreatic cancer xenograft gene expression analysis

To illustrate the value of SMAP, we used a cohort of pancreatic adenocarcinoma (PDAC) patient-derived xenografts ([13]). PDAC is a neoplastic disease with one of the highest proportions of tumor associated stroma which is increasingly recognized as having a major clinical impact. The cohort is composed of 30 human PDAC grafted in immunocompromised mice.

Using SMAP, we separated the tumor and stromal compartments, and retrieved the pathways specific to each of them (figure 5, full table in Supplementary figure 2). The heatmap of human-mouse orthologs reveals specific pathway activity between the tumoral amd stromal compartments despite a high homology rate of the ortologs involved. The heatmap clearly shows the absence of expression of genes involved in biological functions generally attributed to transformed epithelial tumor cells in the stromal compartment such as cytokeratins, proliferation (*e.g*. Cell cycle, DNA replication), DNA damage repair (*e.g*. Homologous recombination, mismatch repair) or more specifically to PDAC (*e.g.* vitamin digestion and absorption, metabolic functions). Conversely, stromal-specific pathways such as those expressed by specific immune cell population (*e.g.* Natural killer cytotoxicity, B-cell receptor pathway) or cancer-associated fibroblasts (*e.g.* related to smooth muscle and extracellular matrix or ECM) are expressed by murine-stromal cells. Well-known oncogenes that are highly homologous between human and mouse were found as over-expressed in the tumoral compartment compared to the stroma (Supplementary figure 3).

**Figure 4.**
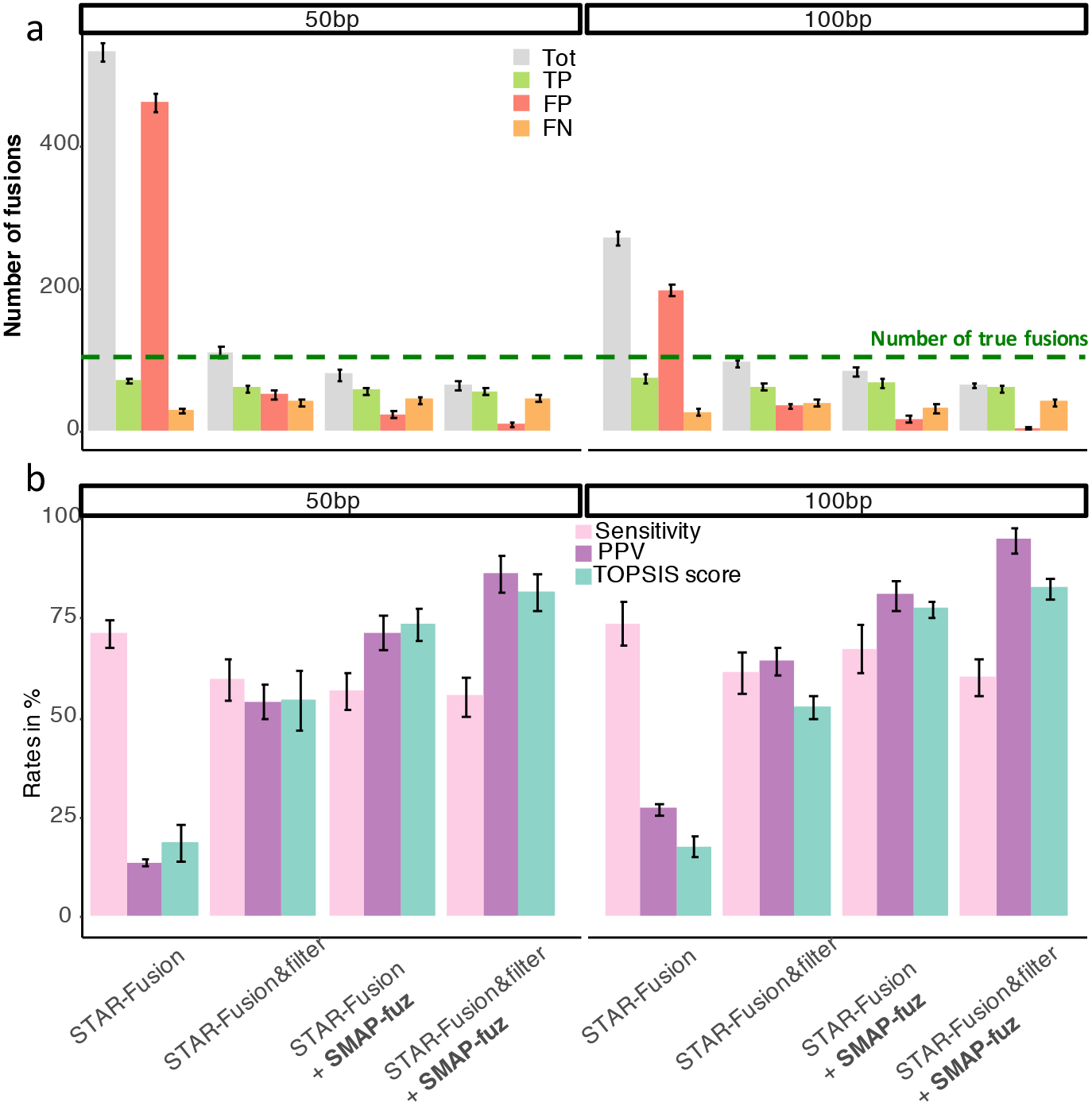
Evaluation of SMAP-fuz on the 100 simulated fusions. **a.** Number of total fusions (Tot), True Positive (TP), False Positive (FP) and False Negative (FN) fusions for each method using 50bp or l00bp sequencing reads. **b.** Evaluation metrics of the different approaches for fusions detections. Bars represent the mean and error bars the standard deviation of five replicate simulation experiments.

To validate the specificity of the stromal and tumor gene expression profiles generated by SMAP, we sought to estimate the relative quantities of cell populations generally found in the tumor microenvironment using MCPcounter [14]. As shown in figure 5b, nearly all cell types are exclusively expressed by the stromal-murine cells, especially fibroblasts as expected.

Finally, to illustrate the value of SMAP fusion, we report (figure 5c) the number of false fusions (human-mouse fusion transcripts) after the application of the SMAP-fuz filter in a 10-fold cross-validation setting (i.e. learning HO parameters on 90% of the total fusion and applying the trained filter on remaining unseen 10%). The results show a significant decrease in the number of false fusions reported by SMAP-fuz in the PDAC cohort.

**Figure 5.**
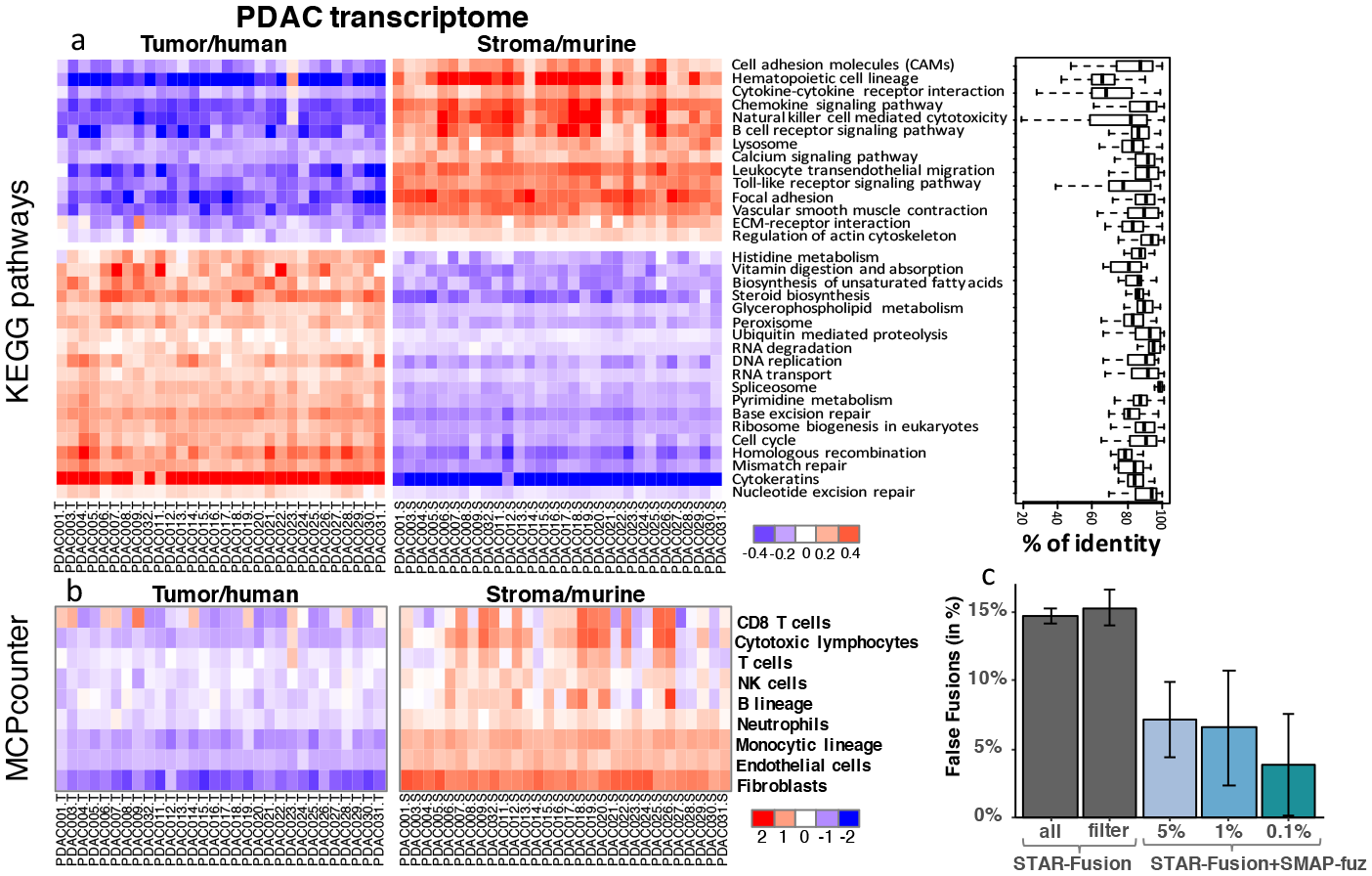
Pathway enrichment analysis. **a.** Pathway enrichment analysis of the human tumor and murine stromal transcriptomes using GSEA. Representation of the metagene expression from the top 20 significant pathways in both direction. Box plots represent the variability of sequence identity percentages between human and mouse orthologs for each pathway. **b**. MCPcounter estimations of tumor microenvironment cellular populations quantification. **c**. Proportions of False Fusions, determined as inter-species chimeric transcripts, determined by different filtering curation approaches, including SMAP-fuz. These rates were computed in a cross-validation setting in which SMAP-fuz null distributions are estimated on a subpart of the samples and used to filter fusions on the rest of the series.

## Discussion and conclusion

As PDX is a method of choice for in vivo drug discovery its specificity as a hybrid species system requires dedicated sequencing processing approaches. Indeed, we show in this study that naive approaches, consisting in ignoring that PDX tumors are mixed cells of different species (noted independent), can result in up to 10% overall misassigned reads and 30% in genes with high homology. The effect of spurious false positive species-assigned reads may have significant effects on any downstream analyses, for instance in variant calling where all human/murine divergence may be considered as a mutation. As well, for gene expression, in particular in cases in which the murine gene is highly expressed as compared to its human ortholog. Therefore, ill-advised processing of PDX sequencing data can introduce a large bias to subsequent analyses.

In order to overcome such consequential issues, we propose a simple and effective approach termed SMAP for simultaneous mapping. We show the superiority of our proposed approach over the few available ones. SMAP only implies minor modifications of standard sequencing processing pipelines and results in major improvements. In particular, SMAP increases the yield of correctly mapped reads while preserving a virtually null level of false positive species assignments. We also show that high homologies between host and graft genomes have virtually no impact on the accuracy of read alignment in terms of species-assignment when using SMAP. Moreover, the computation time necessary for SMAP to process mixed raw reads into two complete transcriptomes is inferior to other methods (see Supplementary figure 4).

In addition to improving xenografts sequencing, we also propose to take advantage of xenografts to increase the reliability of available fusion detection algorithms. The approach, termed SMAP-fuz, uses the spurious human-murine fusions to build a null model of false positives and better retrieves fusion transcripts that stand above the background noise. This strategy aims to be a replacement for *filter*- and *threshold-based* approaches by rather adapting to specificities of each sequencing dataset and fitting null model in a data-driven approach. We showed in both simulated and real datasets that this approach effectively removes false positive fusions making the detected sets of chimeric transcripts more reliable. In its current usage, SMAP-fuz uses two input metrics to estimate the background distribution, the number of spanning fragments and the number of junction reads. However, as additional characteristics are developed to score putative fusion transcripts, for instance expression consistency or sequence homology, these can be simply integrated into SMAP-fuz null model to further improve the identification of chimeric genes.

In this work, SMAP was assessed on simulated transcriptomes of human-murine xenografts as well as on real pancreatic PDX. The use of a real dataset shows the extent to which the complex species mixture can be highly valuable to computationally reconstruct the transcriptome of both the tumor and stromal compartments.

## Materials and methods

### PDX Transcriptome simulations

Xenograft transcriptomes were simulated in silico as a mix of half human and half murine pair-end sequencing reads simulated from a subset of ortholog genes. All human/murine orthologous gene pairs were downloaded from the MGI (Mouse Genome Informatics: www.informatics.jax.org). All pairs of Human/Murine transcripts were blasted and those with a significant match of at least 1kb were kept. From these selected pairs, five sets of 50 homologous transcript pairs were randomly selected based on their level of homology from highly homologous (more than 95%) to low homology (less than 80%).

In order to rigorously control the levels of sequencing error and mutation on read simulations, reads were simulated using wgsim (github.com/lh3/wgsim, [15]) based on the ensembl release 75 cDNA reference of the human (GRCh37) and murine (GRCm38) transcriptomes. 10,000 pair-end reads were simulated for each set of the selected human and murine ortholog pairs. Whole transcriptomes, i.e. using all murine and human transcripts, were simulated using different settings of sequencing errors and mutations as listed and named in table 1. The entire simulation procedure was repeated 5 times. As wgsim is originally proposed as a DNA read simulator which is used here on a reference transcriptome, we investigated whether the use of an RNA-specific read simulator would impact the results and our conclusion. To do so, we simulated reads using Polyester [16] using the same settings. For each setting, 5 classes of expression (10 genes in each) were considered: 5, 25, 70, 200 or 700 reads on average. The entire simulation procedure was repeated 5 times and the results reported in supplementary figure 5 shows that using DNA or RNA specific reads simulators has no impact on this comparative study.

### Fusion simulations

Fusim [17] was used to simulate 100 fusion transcripts from the Human reference transcriptome (Homo sapiens GRCh37 build, ensembl v75). RNAseq data was simulated using wgsim with 50M reads from the total human transcriptome, 20M reads from the total mouse transcriptome and 500 000 from the 100 human fusion transcripts. Fusion transcripts were simulated with diversity of expression level as shown by a high range of RPKM (Supplementary figure 6). Reads were simulated using the *Mutations and errors* settings (table 1). Simulations with 50 and 100 read-lengths were performed. The entire simulation procedure, including the simulation of a novel fusion transcript, was repeated 5 times.

### Species-assignment and fusion evaluation metrics

Identical sets of raw read pairs (in FASTQ format) were processed by each tested pipelines. To quantify the number of correctly species-assigned reads, only uniquely mapped and properly paired reads were kept. STAR-Fusion (STAR-Fusion: Fast and Accurate Fusion Transcript Detection from RNA-Seq. bioRxiv) was applied for fusion transcript detection with. STAR-fusion proposes its own filtering process based on blast analyses (see https://github.com/STAR-Fusion/STAR-Fusion/wiki).

Evaluation metrics were estimated as follow:

TP: True positive. Correctly identified fusions.
FP: Identification of false fusions
TF: Total number of fusions
Sensitivity (%) = (TP/TF) * 100
PPV (Positive predictive value in %) = (TP/(TP+FP)) * 100

TOPSIS (Technique for Order of Preference by Similarity to Ideal Solution [18]) analysis was performed to make decisions on the basis of multiple criteria results. For each approach, TOPSIS scores were calculated by equally weighting the two criteria: sensitivity and PPV.

### Pancreatic Patient-Derived Xenograft dataset and analysis

The mRNA sequencing transcriptome of 30 pancreatic Adenocarinoma PDX ([13]) Pancreatic adenocarcinoma therapeutic targets revealed by tumor-stroma cross-talk analyses in patient-derived xenografts. Cell Reports) were taken from accession E-MTAB-5039. RNA sequencing was Mapped using SMAP with STAR as mapper on ensembl (release 75) with feature count for gene counting and upper quartile normalization was applied. Expression matrix was retrieved only for genes with a unique ortholog between human and mouse (13,708). Mouse and human expression data were scaled for direct comparative analysis.

Filtering for invariant genes: the mean absolute deviation (MAD) was calculated for each gene and those on the first quartile (25%) were filtered out (10,281 genes remaining). GAGE R package [19] was used to perform Gene Set Enrichment Analysis (GSEA) analysis on the scaled gene expression dataset after sample-based zero mean and unit variance scaling. Gene sets were built based on KEGG database. The percentages of sequence identity between human and mouse orthologuous genes where calculated using BLAST and considering human gene sequences as the reference.

### SMAP implementation and usage

SMAP is freely available on github (cit-bioinfo.github.io/SMAP/) and contains python, bash and R scripts. The SMAP pipeline differs slightly from a normal sequencing analysis pipeline by changing the reference genome/transcriptome and by adding one additional post-mapping processing step to separate human and mouse reads.

After downloading the host and graft species genomes (and transcriptomes for RNAseq), these are processed by the `SMAP_prepareReference.sh` shell script which concatenates the genomes. The new reference genomes are also modified to keep track of the species of origin. A standard alignment step can then be applied to the raw xenografts sequences. The SMAP pipeline can then be used to obtain a gene-count matrix for each species (*i.e.* for RNAseq) or to split the aligned reads in BAM format to obtain one BAM per species for subsequent analysis (*e.g.* mutation calling, copy number analysis). Additionally, a module integrated in the SMAP pipeline named SMAP-fuz takes as input STAR-Fusion (initiated by Stransky *et al.* [20] and implemented in Haas et al. (STAR-Fusion: Fast and Accurate Fusion Transcript Detection from RNA-Seq. bioRxiv). available at github.com/STAR-Fusion/STAR-Fusion/releases) output to filter out the spurious fusions.

**List of abbreviations**
SMAP: Simultaneous Mapping
PDX: patient derived xenograft
PDAC: pancreatic adenocarcinoma
PPV: positive predictive value
ECM: extracellular matrix
MGI: Mouse Genome Informatics
RPKM: Reads Per Kilobase Million
TOPSIS: Technique for Order of Preference by Similarity to Ideal Solution
TP: true positive
FP: False positive
TF: Total number of fusions
MAD: mean absolute deviation
GSEA: Gene Set Enrichment Analysis

## Declarations

### Ethics approval and consent to participate

Not applicable.

### Consent for publication

Not applicable.

### Availability of data and material

Not applicable.

### Authors’ Contributions

YB and RN have contributed equally to the development of the methodology and the analyses. YB, RN, AdR, LM, ND and JI participated in writing of the final manuscript. All authors read and approved the final manuscript.

## Acknowledgements

Not applicable.

## References

1. Gao, H., Korn, J.M., Ferretti, S., et al.: High-throughput screening using patient-derived tumor xenografts to predict clinical trial drug response. Nature Medicine 21(11), 1318–1325 (2015)

2. Townsend, E.C., Murakami, M.A., Christodoulou, A., et al.: The Public Repository of Xenografts Enables Discovery and Randomized Phase II-like Trials in Mice. Cancer Cell 29(4), 574–586 (2016)

3. Isella, C., Terrasi, A., Bellomo, S.E., Petti, C., Galatola, G., Muratore, A., Mellano, A., Senetta, R., Cassenti, A., Sonetto, C., et al.: Stromal contribution to the colorectal cancer transcriptome. Nature genetics 47(4), 312–319 (2015)

4. Joyce, J.A., Pollard, J.W.: Microenvironmental regulation of metastasis. Nature reviews. Cancer 9(4), 239–252 (2008)

5. Junttila, M.R., de Sauvage, F.J.: Influence of tumour micro-environment heterogeneity on therapeutic response. Nature 501(7467), 346–354 (2013)

6. Tape, C.J., Ling, S., Dimitriadi, M., McMahon, K.M., Worboys, J.D., Leong, H.S., Norrie, I.C., Miller, C.J., Poulogiannis, G., Lauffenburger, D.A., Jørgensen, C.: Oncogenic KRAS Regulates Tumor Cell Signaling via Stromal Reciprocation. Cell 165(4), 910–920 (2016)

7. Conway, T., Wazny, J., Bromage, A., Tymms, M., Sooraj, D., Williams, E.D., Beresford-Smith, B.: Xenome–a tool for classifying reads from xenograft samples. Bioinformatics (Oxford, England) 28(12), 172–8 (2012)

8. Chivukula, I.V., Ramsköld, D., Storvall, H., Anderberg, C., Jin, S., Mamaeva, V., Sahlgren, C., Pietras, K., Sandberg, R., Lendahl, U.: Decoding breast cancer tissue–stroma interactions using species-specific sequencing. Breast Cancer Research 17(1), 109 (2015)

9. Ahdesmäki, M.J., Gray, S.R., Johnson, J.H., Lai, Z.: Disambiguate: an open-source application for disambiguating two species in next generation sequencing data from grafted samples. F1000Research 5 (2016)

10. Liu, S., Tsai, W.-H., Ding, Y., Chen, R., Fang, Z., Huo, Z., Kim, S., Ma, T., Chang, T.-Y., Priedigkeit, N.M., et al.: Comprehensive evaluation of fusion transcript detection algorithms and a meta-caller to combine top performing methods in paired-end rna-seq data. Nucleic acids research 44(5), 47–47 (2016)

11. Kumar, S., Vo, A.D., Qin, F., Li, H.: Comparative assessment of methods for the fusion transcripts detection from rna-seq data. Scientific reports 6, 21597 (2016)

12. Fisher, S. Ronald Aylmer: Statistical Methods for Research Workers, 11th ed.(rev.) edn. Edinburgh : Oliver and Boyd, ??? (1950). Includes bibliographical references (p. 336–350) and index

13. Nicolle, R., Blum, Y., Marisa, L., Loncle, C., Gayet, O., Moutardier, V., Turrini, O., Giovannini, M., Bian, B., Bigonnet, M., et al.: Pancreatic adenocarcinoma therapeutic targets revealed by tumor-stroma cross-talk analyses in patient-derived xenografts. Cell reports 21(9), 2458–2470 (2017)

14. Becht, E., Giraldo, N.A., Lacroix, L., Buttard, B., Elarouci, N., Petitprez, F., Selves, J., Laurent-Puig, P., Sautès-Fridman, C., Fridman, W.H., de Reyniès, A.: Estimating the population abundance of tissue-infiltrating immune and stromal cell populations using gene expression. Genome Biology 17(1), 218 (2016)

15. Escalona, M., Rocha, S., Posada, D.: A comparison of tools for the simulation of genomic next-generation sequencing data. Nature Reviews Genetics (2016)

16. Frazee, A.C., Jaffe, A.E., Langmead, B., Leek, J.T.: Polyester: simulating rna-seq datasets with differential transcript expression. Bioinformatics 31(17), 2778–2784 (2015)

17. Bruno, A.E., Miecznikowski, J.C., Qin, M., Wang, J., Liu, S.: FUSIM: a software tool for simulating fusion transcripts. BMC Bioinformatics 14, 13 (2013)

18. Hwang, C.L., Lai, Y.J., Liu, T.Y.: A new approach for multiple objective decision making. Computers & operations research (1993)

19. Luo, W., Friedman, M.S., Shedden, K., Hankenson, K.D., Woolf, P.J.: GAGE: generally applicable gene set enrichment for pathway analysis. BMC Bioinformatics 10(1), 161 (2009)

20. Stransky, N., Cerami, E., Schalm, S., Kim, J.L., Lengauer, C.: The landscape of kinase fusions in cancer. Nature communications 5, 4846 (2014)

